# Social association predicts immunological similarity in rewilded mice

**DOI:** 10.1101/2023.03.15.532825

**Authors:** A. E. Downie, O. Oyesola, R. S. Barre, Q. Caudron, Y.-H. Chen, E. J. Dennis, R. Garnier, K. Kiwanuka, A. Menezes, D. J. Navarrete, O. Mondragón-Palomino, J. B. Saunders, C. K. Tokita, K. Zaldana, K. Cadwell, P. Loke, A. L. Graham

## Abstract

Environmental influences on immune phenotypes are well-documented, but our understanding of which elements of the environment affect immune systems, and how, remains vague. Behaviors, including socializing with others, are central to an individual’s interaction with its environment. We tracked behavior of rewilded laboratory mice of three inbred strains in outdoor enclosures and examined contributions of behavior, including social associations, to immune phenotypes. We found that the more associated two individuals were, the more similar their immune phenotypes were. Social association was particularly predictive of similar memory T and B cell profiles and was more influential than sibling relationships or worm infection status. These results highlight the importance of social networks for immune phenotype and reveal important immunological correlates of social life.

---

One of the fundamental roles of an organism’s immune system is to mediate its interaction with its environment (*1*). Immune phenotypes of humans and other species exhibit considerable non-heritable, environmentally-derived variation (*2*–*4*). For example, non-heritable variation in abundance of many types of T and B cells in humans is >80% (*5*). Uncovering which elements of the environment contribute to this variation, and how they do so, remains an open challenge, crucial both for medical practice (*3*) and for understanding the evolutionary and ecological forces that shape the immune system (*6*).

A key part of an individual’s environmental interface is whom they interact with and how often – their social network. Social networks can shape the transmission and exchange of microbes, whether pathogenic (*7*) or non-pathogenic: for example, in the wild, microbiome similarity between individuals correlates with social group (*8, 9*) and the strength of their social ties (*10, 11*). Microbial exposure like this strongly influences immune phenotype; exposing lab mice to various symbiotic microbes shapes their immune phenotypes (*12*–*14*), while systematic enrichment of microbiota produces immune phenotypes quite distinct from standard specific-pathogen free lab mice (*15*–*19*). Individuals who co-habit while co-parenting a child are more immunologically similar to each other than they are to other individuals (*20*). Thus, social interactions could be an important influence on immune phenotype. Ties between elements of social behavior and immune phenotype have been previously identified – for example, both IFN- γ and TNF-α levels have been associated with gregariousness (*21, 22*). But it is unclear how an individual’s immune phenotype is shaped by social life, especially the frequency of interactions and features of the interacting partner(s).

We hypothesize that individuals with stronger social connections should have more similar immune phenotypes. We tested this hypothesis using “rewilded” laboratory mice that are born indoors and then released into outdoor enclosures, where they experience natural weather conditions, eat a varied diet, and have space to roam and burrow. Such settings offer insight into environmental drivers of variation in immune function (*23*). Predators are excluded and chow and water are provided. We used three founder strains of the Collaborative Cross with documented differences in behavior in the lab (*24*): C57BL/6J, 129S1/SvImJ and PWK/PhJ.

Each enclosure contained mice from only one strain; we repeated the experiment while rotating strain-enclosure pairings. We tracked behavior with subcutaneous radio-frequency identification (RFID) tags; five RFID stations per 180 m^2^ enclosure – one at the chow feeder and four arrayed in a diamond pattern around the perimeter (Fig. 1A) – recorded visits by each mouse during each five-week experimental period. We collected blood samples for Complete Blood Count (CBC) analysis prior to release and two weeks post-release, when we challenged a subset of mice with *Trichuris muris*, an intestinal nematode parasite. At five weeks post-release, prior to any shedding of *T. muris* eggs, we trapped out the mice for extensive immune phenotyping (*25*) and collected fecal samples for 16S microbiome analysis. We analyzed our data using Bayesian linear regression models with appropriate response variable distributions and priors to generate posterior probability distributions for the associations between predictors and response variables (see Methods).

**Figure 1:**
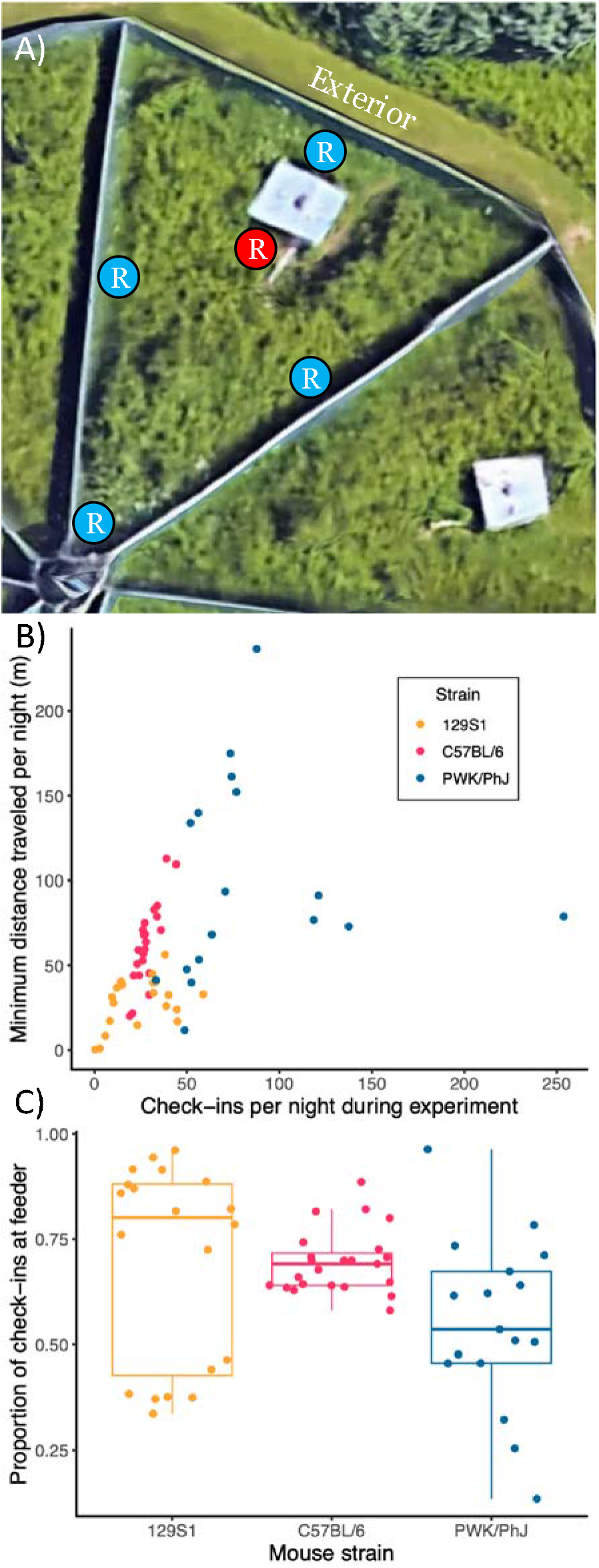
Rewilded mouse activity levels vary within and among strains. A) Aerial image via Google Earth of one of the three enclosures used during the experiment (here, enclosure #4). “R” circles identify the locations of RFID stations within enclosure (red = feeder; blue = non-feeder); the reader layout was the same for each enclosure. B) Check-ins and minimum distance traveled per night for each individual. C) Proportion of check-ins taking place at the RFID reader attached to the feeding station within each enclosure, for each individual.

Individual behavior, in terms of both abundance and spatial and temporal distribution of check-ins at RFID stations, varied substantially both within and among strains. PWK/PhJ mice (*n* = 17) had the most check-ins, followed by C57BL/6 mice (*n* = 23) and then 129S1 mice (*n* = 20) (Fig. 1B, Table S1). PWK/PhJ and C57BL/6 mice traveled similar minimum distances per night but generally further than 129S1 mice (Fig. 1B, Table S2). Strain did not predict proportion of check-ins occurring at the feeding station (Table S3), but there was wide variation within strains (Fig. 1C). In general, strain exerted some influence on rewilded mouse behavior, but there was substantial additional variation not accounted for by genetic background.

To study social behavior, we calculated social networks for each co-housed group based on overlapping appearances at RFID stations. We defined pairwise association strength (*n* = 362 pairs) with the simple ratio index (SRI), here the ratio of the number of night-location combinations at which the two mice appeared within some time interval of each other to the total number of night-location combinations at which one or both mice appeared (*10, 26*). Much like individual activity, observed social associations varied within and among strains. C57BL/6 mice and PWK/PhJ mice both had stronger pairwise associations over fifteen-minute overlap windows than 129S1 mice (Fig. 2A, Table S4, Fig. S1). PWK/PhJ and C57BL/6 mice had similar average association strengths despite PWK/PhJ mice having many more check-ins than C57BL/6 mice (Fig. 1B). Together these results suggest that levels of activity are not the sole drivers of association between individuals; furthermore, cage-sharing prior to release and sibling relationships did not influence association strengths (Fig. S2, Table S4). Associations were stronger at feeding stations than at non-feeding locations for all strains, but pairs’ associations at the locations were correlated (Pearson’s *r* = 0.524) and mice did associate at non-feeding locations (mean strength of social associations at non-feeders: 0.196) (Fig. 2B). Intriguingly, despite the variation in pairwise association strength, within each network mice were generally quite similar in their average association strength and centrality (Fig. S3). The experimental challenges with *T. muris* had negligible effects on individual and social behavior, with the only small effects being slightly decreased check-in counts and a slight increase in the relative proportion of check-ins at the feeder by helminth-infected mice (Tables S1–S5). Overall, we find substantial genetic differences in individual and social behavior in a semi-natural setting and further within-strain heterogeneity that allows us to examine how social behavior interacts with immune phenotypes.

**Figure 2:**
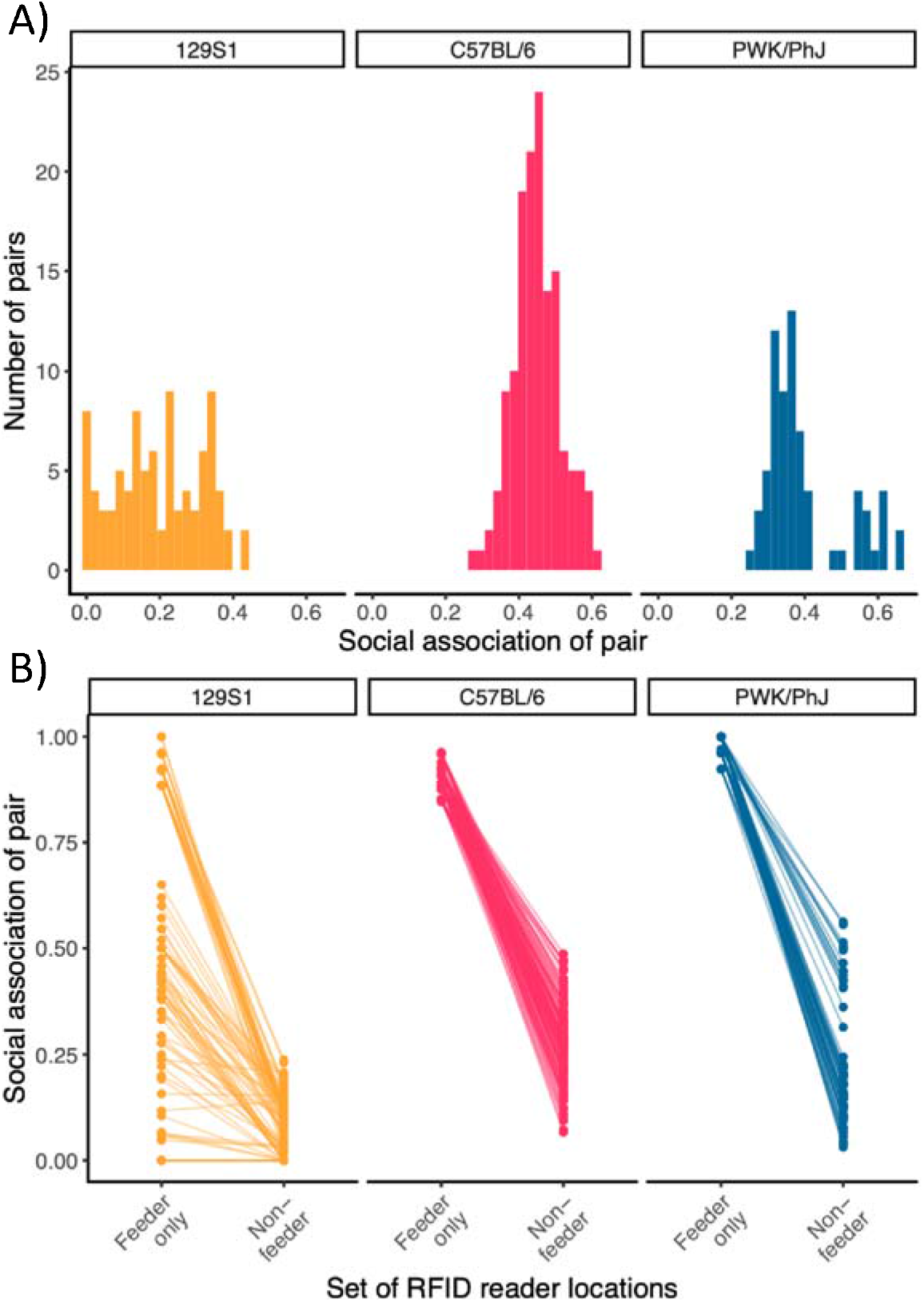
Rewilded mouse pairwise association levels vary within and among strains and locations. Associations for this plot were calculated with a 15-minute overlap threshold. A) Strength of social associations for each pair of mice, broken down by genotype. B) Pairwise social association strengths by set of locations considered: only associations at RFID reader at feeding station vs. only associations at all other RFID readers.

We next turned to assessing our hypothesis that immune phenotype and behavior would be linked. We found that individual-level behavior – e.g. number of check-ins – mostly does not predict immune phenotypes of mice (Table S6). However, we did find extensive evidence that social interactions shaped the immune phenotypes of the mice. We calculated pairwise similarities of several different aspects of immune phenotype at the time of trapout: white blood cell profiles drawn from CBC measurements, CD4, CD8, and combined CD4 and CD8 T cell memory phenotypes (determined from cell surface expression of CD44 and CD62L as measured by flow cytometry) in blood and mesenteric lymph nodes (MLNs), B cell phenotypes in the MLNs drawn from flow cytometry, plasma cytokine concentrations, and MLN cytokine production from antigenic stimulation (*25*). To quantify similarity, we used Jaccard index for cell type distributions and Manhattan distance for cytokine measures. We found that strength of social association of a pair correlated positively with pairwise similarity of several aspects of immune phenotype: most strongly with CD4 T cell memory phenotypes in the MLNs (*n* = 362 dyads), but combined MLN CD4/CD8 T cell memory (*n* = 362); MLN B cell phenotypes (*n* = 362), and white blood cell profiles from CBC differentials (*n* = 391) also exhibited positive correlations between social association and immune similarity (Fig. 3A, 3B, S4, Table S7).

**Figure 3:**
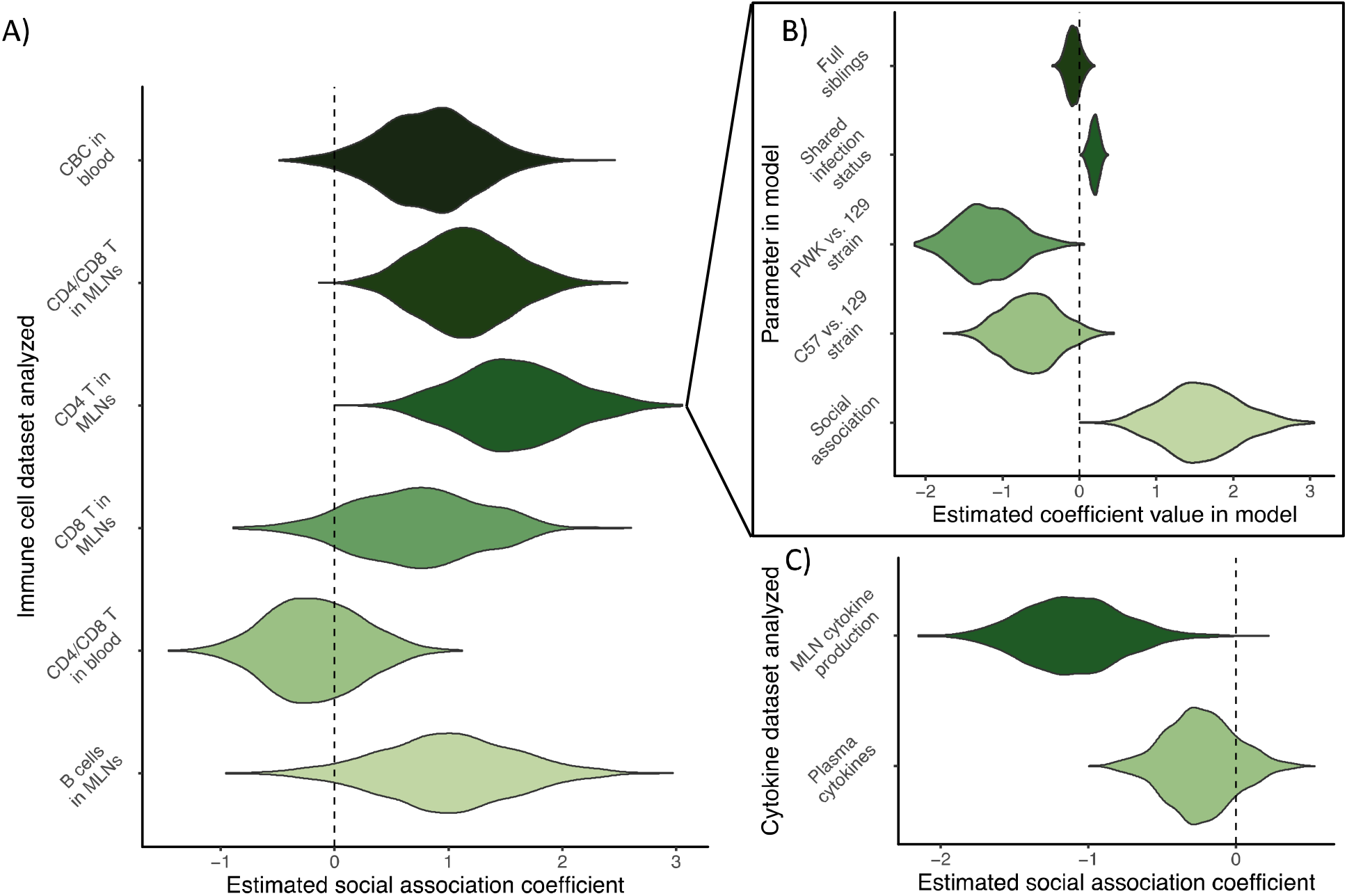
Social association correlates with immune similarity for several aspects of immune phenotype. Violin plots show regression model coefficient posterior probability distributions plotted via 1000 samples from model-estimated parameter value distribution. A) Posterior probability distributions for relationship between social association and cellular immune similarity, estimated by Bayesian linear models. Other predictor variables in models are strain of dyad, shared infection status, and sibling relationships; individual identity is included as a random effect. B) Full model results for fixed-effect predictors from model of CD4 T cell memory phenotype similarity in mesenteric lymph nodes (MLNs). C) Posterior probability distributions for relationship between social association and similarity of aspects of cytokine phenotype.

Thus, these results indicate a form of social network assortativity (*27*): mice that associated more had more similar immune phenotypes. We did not find such relationships for blood T cells (*n* = 323), or for plasma cytokines (*n* = 306); intriguingly, *in vitro* MLN cell cytokine production in response to stimulation showed the opposite relationship (*n* = 147) (Fig. 3A, 3C, Tables S7, S8). In addition to effects from social association, we consistently found that different strains exhibited different levels of immune variability; shared infection status usually had a small positive effect on immune similarity, while sibling relationships consistently had none (Fig. 3B, Tables S7, S8). Overall, these results suggest that social association can predict similarity of WBC differentials in the blood and memory lymphocyte composition in lymphoid tissue.

We next used CBC data from blood draws prior to release and at two and five weeks after release to examine how the relationship of immune similarity to social association changed across each experiment. We found that CBC similarity prior to release did not correlate with social association during the experiment and that a weak potential correlation emerged between CBC similarity at the midpoint and social association during the experiment (Fig. 4A, Table S9). These results contrast with the appreciable relationship at the end of each block. And immune similarity prior to release only weakly correlated with immune similarity at the end (Pearson’s *r* = 0.202), with greater CBC variation on average at the end of each block (Fig. S5). Accordingly, we find that the relationship between immune similarity and social association is not coincidental; rather, it emerges during the experiment, and perhaps the social associations are structuring the immunological changes developing during the experiment.

**Figure 4:**
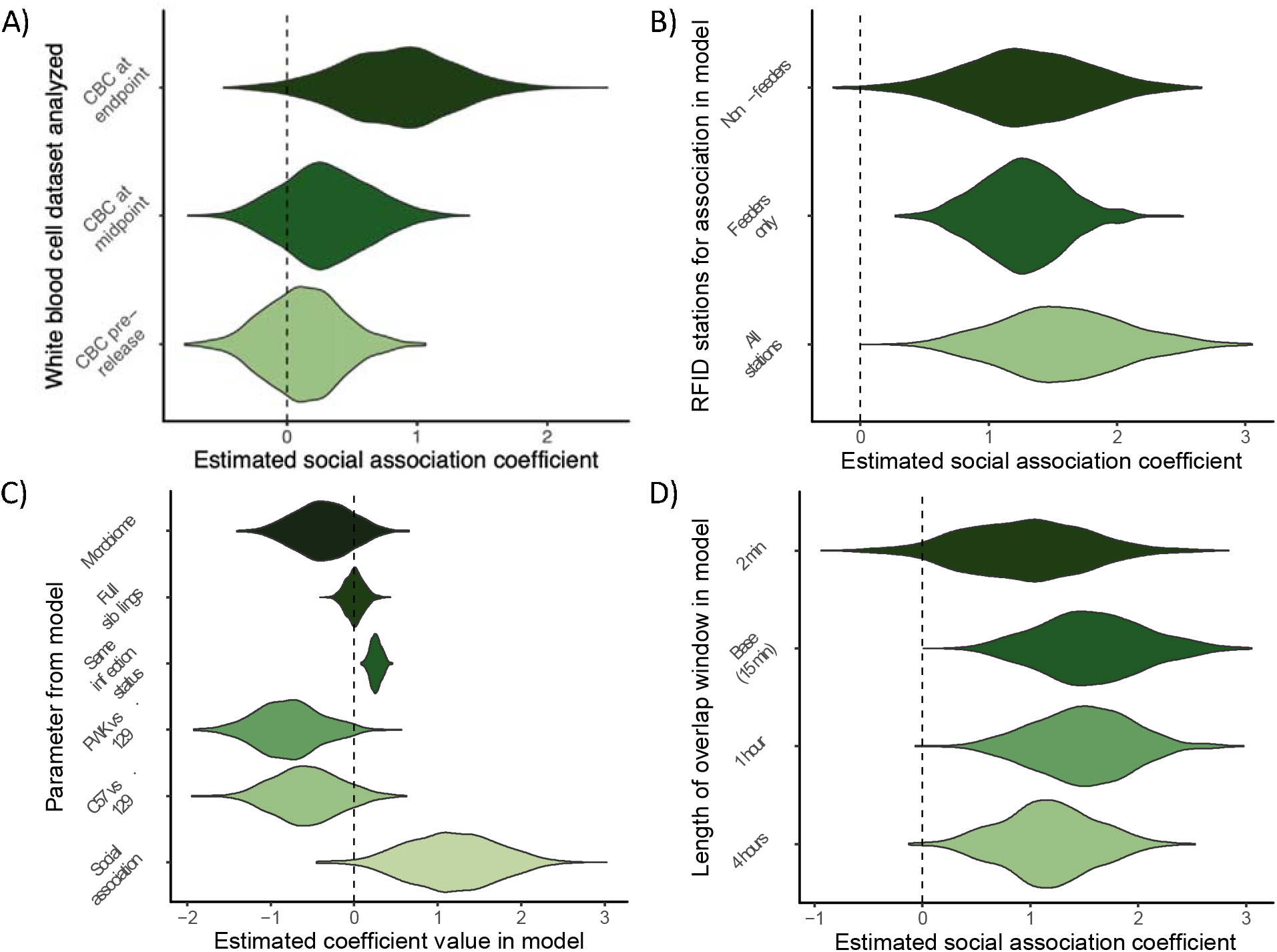
Exploring hypotheses for the sociality-immunity link. Violin plots show regression model coefficient posterior probability distributions plotted via 1000 samples from model-estimated parameter value distribution. A) Estimated relationships between social association during experiment and similarity of white blood cell profiles (via CBC profiling) at three different timepoints during experiment. B) Estimated relationships between social association calculated from different subsets of RFID readers and similarity of MLN CD4 T cell memory phenotypes. C) Full results for fixed-effect predictors from model of MLN CD4 T cell memory phenotype similarity in a model that includes gut microbiome similarity. Note that model selection methods do not prefer a model with microbiome similarity over one without. D) Estimated relationships between social association calculated from different overlap window lengths and MLN CD4 T cell memory phenotype similarity.

Using different metrics for association and environmental co-variates, we can investigate possible mechanisms for such a relationship: direct microbial transmission, similar space use patterns (*28*) and potentially therefore environmental acquisition of microbes (*29*), and/or shared dietary proclivities (*30, 31*). Since diet can influence immune phenotypes (*30, 31*), shared diets could drive immunological similarity. If so, social association at the feeder station may be relatively more predictive of immune similarity than social association at the non-feeder stations. We calculated association strengths at just these location subsets for each pair and assessed their correlations with CBC and MLN CD4 T cell memory similarity. We found that non-feeder and feeder associations predicted CD4 T cell memory similarity approximately equally well (Fig. 4B, Table S10), while non-feeder associations predicted CBC similarity better than did feeder associations (Fig. S6A, Table S11). These results suggest that diet similarity is not a key driver of immune similarity, although more detailed analysis of diet – for example, through metabarcoding – will be necessary to investigate this possibility further.

Another possible explanation is that social associations lead to microbe transmission through direct contact and/or indirect, environmentally-mediated transmission, with these shared microbes driving immunological similarity. We found no evidence of a range of common mouse pathogens circulating in our rewilded mice (Table S12), suggesting that disease transmission, which could explain this observed relationship, is not taking place. We analyzed fecal microbiome samples from the end of the experiment and examined the relationship between gut microbiome similarity, social association, and immune similarity. We did not find evidence of a relationship between microbiome similarity and immune similarity for CD4 T cell memory phenotype in the MLNs (*n* = 289) (Fig. 4C), CBC phenotype (*n* = 315) (Fig. S6B), or B cell phenotype in the MLNs (*n* = 289) (Table S13). Our model selection methods preferred models of immune similarity excluding microbiome similarity, and social association patterns did not positively correlate with gut microbiome similarity (*n* = 338) (Table S13). These results suggest that the gut microbiome at experimental endpoints is not driving the observed relationship between social interactions and immune phenotypes; however, it does not rule out the potential influence of cumulative exposures to microbes or to microbial antigens.

We next looked at predictive power for social association defined from longer and shorter overlap windows. Associations calculated from longer overlap windows, because individuals do not have to be present at a location as close in time to each other and are therefore less likely to have physically interacted, are weaker indicators of direct contact, and therefore direct microbe transmission, between individuals. But they do still indicate shared space use patterns, which may mean similar environmental microbe acquisition (as well as opportunities for indirect microbe transmission). If associations from shorter overlap windows are more predictive of immunological similarity, then direct contact may be a key driver of the effect. We calculated social association for four-hour, one-hour, and two-minute overlap windows, in addition to our default fifteen-minute window. We found that all four metrics could predict CD4 T cell memory similarity, with one-hour associations being the most predictive but broadly similar predictive power for all four metrics (Fig. 4D, Table S14). Contrastingly, shorter time intervals may be more predictive of CBC similarity (Fig. S6C, Table S15). These results suggest that shared space use helps to explain immunological similarity – as do the similar results for associations at different sets of RFID readers – but direct contact or traveling together may be more important for some aspects of immune phenotype, such as the myeloid cells (monocytes, neutrophils, etc.) that appear in our CBC dataset but not the flow cytometry dataset.

Taken together, these analyses support our focal hypothesis that social interaction influences immune phenotypes. We are limited in our ability to explain exactly how this influence is exerted. Shared antigenic experience is our most plausible explanation, given the role of social contact in transmitting commensals and parasites that shape immune phenotypes, but it cannot be solidified with our data. A key nuance is that only some aspects of immune phenotype, in particular cellular composition, are influenced by social association. The strong influence of sociality on T and B cell phenotypes makes especial sense, given that these are adaptive immune cells activated by specific antigens. The fact that social association predicts adaptive immune similarity in the MLNs but not the blood may reflect the role of the lymph nodes as sites of that antigen recognition. In our analysis of GxE effects on immune phenotype (*25*) we found that environment had more effect on immune cell composition in the blood than in the MLNs. However, here we are investigating within-strain heterogeneity in a single shared environment, so we may have a different predictor for something that would have been noise under a conventional analysis. A caveat for why endpoint microbiota similarity does not predict immune similarity may be that we only examined bacterial communities. In addition to the potential effects of other environmental antigens (including eukaryotes and viruses), the gut bacterial communities we examined are only a snapshot of the antigenic experience of each individual. Although we lack the data to investigate social interaction content – e.g. dominance, or affiliative behavior – it has been shown to shape immune phenotypes, as in non-human primates where social rank correlates with some aspects of immune gene expression (*32, 33*). This process could explain why more associated mice have less similar *in vitro* cytokine production profiles – perhaps there are dominance hierarchies within the networks driving polarization of cytokine responses between associates despite shared antigenic experiences.

Regardless, these results also offer intriguing insight into the flexibility of the immune system in response to new conditions and experiences. The immunity-sociality relationships like we observe here may also be generating and structuring some of the extensive heterogeneity observed in human immune phenotypes (*2, 3*). And if social association influences immune state, and if immune state can predict functional responses (*3*), then individuals that are more associated should be more similar in their susceptibility to a given parasite challenge, at least in some aspects of the immune response – e.g., memory quality and specificity – if not others – e.g., cytokine responses. Heterogeneities in disease susceptibility have been shown theoretically and empirically to impact infectious disease dynamics and pathogen evolution (*34, 35*). Our work here highlights a way that such heterogeneities might emerge and may therefore identify a phenomenon important not only for hosts but also for pathogens.

Overall, we document extensive behavioral variation in laboratory mice rewilded in outdoor enclosures. We show that interactions between individuals shape immune phenotypes such that the more associated two individuals are, the more similar their immune phenotypes. This effect, which emerges during the experiment, is particularly strong for cellular composition and is weak or even negative for cytokines. These results offer intriguing implications for the generation of natural immune variation and the role of social contact in shaping immune systems, and they highlight important new directions of study for understanding disease susceptibility, infectious disease ecology, and the operation of natural selection on immune phenotypes.

## Acknowledgments

We thank William Craigens, Christina Hansen, and the Graham Lab for invaluable assistance in the field. In addition, we thank Mingming Zhao for help with setting up experiments in the lab and Joseph Devlin and Johnson Randall for help in maintaining the Joes’ Flow App Software. We thank Dr. Elia Tait Wojno for providing the *Trichuris muris* parasite larvae to us. We thank Dr. Aura Raulo, Prof. Sarah Knowles, the Knowles Lab, Prof. Robert Froemke, Prof. Shruti Naik, and Prof. William Gause for comments and suggestions.

## Author Contributions

Conceptualization: AED, OO, KC, PL, ALG. Methodology: AED, OO, QC, YC, EJD, RG, KK, DJN, CKT, OMP, ALG. Investigation: AED, OO, RSB, YC, KK, AM, OMP, KZ, PL, ALG. Data curation and analysis: AED, OO, ALG. Hardware and software: AED, QC, RG, JBS. Writing – original draft: AED, OO, PL, ALG. Writing – review and editing: AED, OO, EJD, AM, CKT, KC, PL, ALG. Visualization: AED, OO, ALG. Supervision: KC, PL, ALG. Funding acquisition: KC, PL, ALG. All authors approved the final manuscript.

## Data and materials availability

All metadata, behavioral data, immunological data, and analysis code will be available at https://github.com/aedownie/rewilding2021. Until the link is active, please e-mail the corresponding authors with data requests, which will be fulfilled. RNA expression data will be deposited in the Gene Expression Omnibus.

## List of Supplementary Materials

Methods

Supplementary Text Figs. S1–S11

Tables S1–S20

