## Supplementary Materials for "Social association predicts immunological similarity in rewilded mice"

Contains:

Materials and Methods

Supplementary Text

Figs. S1–S11

Tables S1–S20

Additional References

**Methods**

Re-wilding

For our re-wilding, we used *n* = 89 female mice from three mouse strains used as founders of the Collaborative Cross: C57BL/6, 129S1, and PWK/PhJ. Genetic variation in these strains spans ~50 million SNPs, a similar order of magnitude to the total variation documented among humans (Link et al. 2018, Lusis et al. 2018). The experiment took place across two consecutive blocks. Before each block, mice were bred at the National Institutes of Health before shipping to Princeton University for acclimation in a dedicated animal facility to temperatures and light cycles characteristic of summer in New Jersey (26 ˚C ± 1 ˚C, and a 15-hour light/9-hour dark cycle). Mice were housed in cages of five prior to release; a subset of mice from each cage were chosen for release, while the others were kept as controls in the warm and humid vivarium (Oyesola et al. *in prep*). For release, the mice were divided into three groups per block, with each group consisting of a single strain. In Block 1 we released *n* = 42 mice (15 PWK/PhJ, 14 C57BL/6, 13 129S1); in Block 2 we released *n* = 47 mice (16 PWK/PhJ, 18 C57BL/6, 13 129S1). Strains were kept separate to avoid any potential for antagonistic interactions and, between blocks, we rotated the enclosure to which each strain was assigned so that enclosure pen and mouse strain were unconfounded.

We placed each group of mice in a set of enclosures located in Princeton, New Jersey during the late spring and summer of 2021. Block 1 ran from May–June 2021, Block 2 from July–August 2021. These enclosures (Fig. S1), described in Leung et al. (2018) and also employed in Lin et al. (2020), Yeung et al. (2020), Chen et al. (*in review*), and Budischak et al. (2018), are triangular wedges, ~180 m^2^ in area, and enclosed by 1 m-high walls of zinc-coated steel which penetrate into the ground ~0.5 m. Natural vegetation grows in the wedge, with the exception of a strip around the perimeter of the enclosures which is kept clean-cut for access, and there is a low wooden hut of 180 cm x 140 cm x 70 cm with a corrugated metal roof that provides some shelter. The exterior walls are topped with an electric fence to exclude any terrestrial predators or other animals, while aerial predators are deterred with the expedient of several fishing lines strung over each enclosure and hung with aluminum pie plates. Several years’ experiments indicate that these measures are effective at both keeping mice in and predators and other animals out, with the exception of small birds that pose no threat to the mice. Within each enclosure the mice can roam and forage freely; each enclosure contains one feeding station near the hut stocked with lab chow (PicoLab Rodent Diet 20) and two water bottles inside the hut. We checked the status of these three times a week, topping up the chow if necessary and completely replacing it each week and refilling the bottles if they were running low.

We tracked mouse behavior via RFID check-ins. Prior to release, we injected each mouse with a subcutaneous RFID tag (8 mm x 1.4 mm FDX-B “Skinny” PIT Tag, Oregon RFID, Portland, OR, USA), patched the injection site with liquid bandage, and allowed the mouse time to heal. Each enclosure contained five RFID readers, of a design described in Budischak et al. (2018). Briefly, each reader consisted of a plastic PVC tube ~45 cm in length around which an antenna, connected to an RFID reader/writer, is wrapped. When a mouse runs through the tube, the RFID reader/writer records the tag number of that mouse and transmits it to a central database that records the tag number, the particular RFID reader, and the date and time of the check-in (see further details below). One of the five RFID readers in each enclosure was attached to the feeding station, such that each time the mouse entered or exited the feeding station it produced a check-in. The other four are arranged in a diamond pattern around the perimeter of the enclosure (Fig. 1A).

Prior to shipping from the NIH to Princeton, blood samples for CBC analysis were collected with cheek bleeds; at Princeton, several days prior to release, fecal samples were collected. At 2 weeks after release, we trapped the mice using Longworth live traps baited with peanut butter and a pellet of lab chow. We placed at least 40 traps in each enclosure, inside and around the hut and in other places throughout the enclosure. We drew further blood samples for CBC analysis and collected fecal samples; we placed blood samples on ice, while fecal samples were immediately placed on dry ice and then frozen at -80 ºC. We caught *n* = 6 mice that had lost their original RFID tags since their release into the enclosures; each was given a new RFID tag, and the injection site was patched with liquid bandage. In addition, we selected a subset of individuals (*n* = 24 for Block 1, *n* = 27 for Block 2) for challenges with *Trichuris muris*, a natural gut helminth of mice. We gave each of these mice 200 embryonated eggs via oral gavage. After sampling and infection challenge, mice were re-released in their home enclosure. We also caught mice that had escaped from their original enclosure into different enclosures; we returned these mice to their original enclosures and repaired the breach to prevent further escapes.

At approximately five weeks after release, we re-trapped all mice using the same method. Again we drew blood samples for CBC analysis with cheek bleeds using a 4 mm Medipoint Golden Rod Lancet Blade (Medipoint NC9922361) and collected fecal samples; then, we anesthetized the mice via isoflurane inhalation and collected the remaining blood with a terminal cardiac puncture for analysis via flow cytometry. We processed the mice for collection of the mesenteric lymph nodes, the cecum, and the GI tract. We placed blood samples again on ice, while we placed fecal samples on dry ice and then froze them at -80 ºC. We dissected the cecum for counts of *T. muris* parasites. In total, at the end of the experiment we re-captured *n* = 72 of the original 89 mice released.

Immunological assessment

We performed a CBC analysis with differentials of the whole blood sample collected into 1.3mL heparin coated tube (Sarstedt Inc, NC9574345) and analyzed using the element HT5, Veterinary Hematology Analyzer (Heska) allowing us to abundance of leukocytes and the relative abundances of five different leukocyte types (neutrophils, lymphocytes, monocytes, eosinophils, and basophils) in the blood. We assessed lymphocyte populations in the peripheral blood and in the mesenteric lymph nodes using flow cytometry.

For Blood, whole blood collected via the cheek bleeds into heparinized tubes (Sarstedt Inc SC MTUBE 1.3ML LI HEP/PK100) were mixed with heparinized blood collected via cardiac puncture method. The combined blood samples were spun for 10 minutes at 1500 rpm and plasma was collected and stored at -80^o^C for further cytokine analysis. The cellular component re-suspended in PBS next underwent a density gradient separation process using the Lymphocyte Separation Media (LSM^TM^ MP Biomedicals, LLC, Irvine, CA) according to manufacturer’s instruction. Isolated PBMCs were washed twice in PBS and then used for downstream spectral cytometric analysis. Single cell suspension from the mesenteric lymph nodes (MLNs) were prepared by mashing the tissues individually through a 70μm cell strainer and washed with RPMI. Cells were then washed with RPMI supplemented with 10% FCS. Live cell numbers were enumerated using the element HT5, Veterinary Hematology Analyzer (Heska).

For Spectral Cytometry, single-cell suspensions prepared from the PBMCs and MLN were washed twice with Flow cytometry buffer (FACs Buffer) and PBS prior to incubating with Live/Dead™ Fixable Blue (ThermoFisher) and Fc Block™ (clone KT1632; BD) for 10 minutes at room temperature. Cocktails of fluorescently conjugated antibodies (Table S16) diluted in FACs Buffer and 10% Brilliant Stain Buffer (BD) were then added directly to cells and incubated for a further 30 minutes at room temperature. For the lymphoid panel, cells were next incubated in eBioscience™ Transcription Factor Fixation and Permeabilization solution (Invitrogen) for 12-18 hours at 4°C and stained with cocktails of fluorescently labeled antibodies against intracellular antigens diluted in Permeabilization Buffer (Invitrogen) for 1 hour at 4°C.

Spectral Unmixing was performed for each experiment using single-strained controls using UltraComp eBeads™ (Invitrogen). Dead cells and doublets were excluded from analysis. All samples were collected on an Aurora™ spectral cytometer (Cytek) and analyzed using the OMIQ software (<https://www.omiq.ai/>), data cleaning and scaling was done using algorithms like FlowCut within the OMIQ software. Sub-sampled cells including 10,000 Live, CD45+ cells were reclustered in an unsupervised version using the Joe’s Flow software, (Github: <https://github.com/niaid/JoesFlow>).

We also assessed cytokine phenotypes through two techniques: plasma cytokine levels at time of sacrifice and production of cytokines by MLN cells in response to antigenic stimulus. Plasma concentrations of IL-5, IL-6, IL-22, IL-17A, IFN-γ, TNFα were measured also measured using the commercially available murine Th cytokine LEGENDplex assay (Biolegend) according to the manufacturer’s instructions. For MLN cytokine production assays, single cell suspension of MLN cells were reconstituted in RPMI at 2 × 10^6^ cells/mL, and 0.1 mL was cultured in 96-well microtiter plates that contained 10^7^ *cf.*u/mL UV-killed microbes, 10^5^ αCD3/CD28 beads (11456F), or lipopolysaccharide (LPS; 100ng/mL) (L2630) or PBS control. The microbes included were: *Bacteroides vulgatus* (ATCC 8482), *Candida albicans* (UC820), *Clostridium perfringens* (NCTC 10240), and *Trichuris muris* excretory-secretory antigens.  Supernatants were collected after 2 days and stored at −80°C. Concentrations of IL-5, IL-6, IL-22, IL-17A, IFN-γ in the supernatants were measured using a commercially available murine Th cytokine LEGENDplex assay (Biolegend) according to the manufacturer’s instructions. To quantify the cytokine production, we divided production in response to antigens/microbes by production with 1x PBS, then took the log_2_ of the divided quantity plus 1 (to account for any samples with no production in response to antigenic stimulation).

Microbiome

DNA for microbiome analysis was isolated from frozen fecal samples collected at the endpoint using the QIAsymphony PowerFecal Pro DNA Kit (938036) with the QIAsymphony SP instrument. We prepared the DNA library using the SMRTbell Express Template Prep Kit 2.0 and Amplicon-Seq library sequencing was performed on the PacBio Sequel II system. Sequencing reads were primer trimmed, filtered, dereplicated and then analyzed using the Divisive Amplicon Denoising Algorithm 2 (DADA2) pipeline (<https://benjjneb.github.io/dada2/tutorial.html>). We performed bacterial taxonomy assignment against the DECIPHER ([http://www2.decipher.codes/Downloads.html](https://gcc02.safelinks.protection.outlook.com/?url=http%3A%2F%2Fwww2.decipher.codes%2FDownloads.html&data=05%7C01%7Coyebola.oyesola%40nih.gov%7Cba7fe293217949c0f8de08da7bc876dd%7C14b77578977342d58507251ca2dc2b06%7C0%7C0%7C637958399763821125%7CUnknown%7CTWFpbGZsb3d8eyJWIjoiMC4wLjAwMDAiLCJQIjoiV2luMzIiLCJBTiI6Ik1haWwiLCJXVCI6Mn0%3D%7C3000%7C%7C%7C&sdata=NAGAeNFxKuDHogjfHzmoMMx%2F5lErO5nEUmzRZnrfKkQ%3D&reserved=0)). and SILVA v138 databases. We generated an Amplicon Sequence Variants (ASVs) table, which was then used for assessment of pairwise microbiome similarity. We scaled read counts for each taxon for each individual by the total number of reads for that individual, producing relative abundance estimates. From these relative abundance distributions we calculated microbiome similarity for pairs of mice with Jaccard index.

Individual behavior assessment

We described mouse individual behavior based on RFID visits logged at the five RFID readers in each enclosure. The RFID reader casing houses an RFID reader/writer board (Priority1 Design) and a [Particle](https://www.particle.io/) Photon Wi-Fi-enabled IoT device. These are powered with an external power supply plugged in to an outlet at the center of the enclosures. Each time a mouse passes through the RFID reader, the RFID reader/writer transmits the RFID signal to the Photon unit. The Photon unit then transmits to a central Particle server a message containing the RFID number, the reader identity, and the time of the check-in, which is accurate to the second. We collect the data from the Particle server via a webhook in our own database. The resulting behavioral dataset consists of 103,367 check-ins across the two blocks, with RFID number, date, time, and location for each check-in. Each reader was associated with a particular location in its enclosure: the feeder, the left and right sides (relative to the exterior wall of the triangle), base (along the exterior wall), and tower (the reader closest to the tip opposite the exterior wall) (Fig. 1A). We converted each RFID number to the ID number of the associated mouse. Some mice were associated with two RFID numbers because they received replacement RFIDs at the midpoint trapping during the experiment, and we synonymized these RFID numbers in our database.

We applied several filtering steps to ensure that the data did not contain accidental biases. We first identified times when individual RFID readers were offline for an extended period of time, due either to power and Wi-Fi outages at the enclosures or to water entering the reader’s casing. If at least one reader was offline during a particular night, then we dropped from the dataset all check-ins from all readers within the same enclosure as the affected reader for the corresponding 24-hour period (12 noon to 12 noon), to avoid mis-representing spatial location of activity, relative abundance of activity, or social network structure. For our second filtering step we removed nights during which trapping took place from our check-in dataset, since the number of check-ins for an individual on a given night would depend on when, or if, that mouse entered a trap. In our third step, for each individual that escaped its home enclosure, we also removed from our dataset all of its check-ins taking place outside of the enclosure in which it was initially released. After completion of this filtering, the dataset comprised 97,358 check-ins.

We calculated activity levels using multiple metrics. Because the length of the experiment, in terms of nights of activity, varied from enclosure to enclosure, and because some nights had to be removed due to missing reader data, we calculated all activity levels on a per-night basis. In Block 1, Wedges 2 and 3 have 33 nights of data, while Wedge 4 has 25 nights of data; in Block 2, Wedges 2 and 3 have 26 nights of data and Wedge 4 has 27 nights of data (there are fewer nights for Block 2 due to a power and Wifi outage that delayed the start of tracking until after release). The simplest metric was the average number of check-ins per night for each mouse. We also calculated average minimum distance traveled and roaming entropy each night. Minimum distance was calculated based on the distances between each RFID reader within a given enclosure. Each time a mouse visited a new reader on a given night, the distance between the new reader and the previous reader was added to the distance that mouse had traveled that night. If the reader visited was the same as the previous reader, then no distance was added. Distances between reader locations varied slightly among enclosures due to differences in overall dimensions. An average across all nights was calculated for each mouse to give the minimum distance traveled per night by that mouse.

Roaming entropy describes how evenly an individual’s activity is spread in time and space (Freund et al. 2013). We calculated roaming entropy after the method defined in Freund et al. (2013), using 12-hour windows, from 8 PM to 8 AM, corresponding to the hours of greatest mouse activity (Fig. S9). The roaming entropy *RE_i,t_* for mouse *i* on day *t* is

${RE}_{i,t}=-\sum_{j=1}^{k} p_{i,j,t}\log\left( p_{i,j,t} \right)/log(k)$,

where the reader is *j* and the total number of readers in the wedge is *k*. *p_i,j,t_* is the probability of mouse *i* being observed at reader *j* on day *t*. To calculate this value, we divided each individual night was divided into minute-long windows. For each mouse, the reader most recently visited at the end of that window was assigned as the reader at which the mouse was present during that window (thus assuming that each mouse is in the vicinity of the last reader it visited until it visits a new reader). *RE_i,t_* is on the interval [0,1], with lower values indicating that activity is mostly concentrated at one location and higher values indicating that activity is spread evenly in time around several locations. Mean roaming entropy for an individual was calculated by as the average of their roaming entropies from all nights during which they appeared.

Social behavior assessment

We similarly assessed social behavior using RFID check-in records. We derived our approach and code from that in Raulo et al. (2021), with suitable modifications for our particular experimental set-up. The basic metric we used to calculate pairwise social association strength between individuals is the simple ratio index (SRI). Mice are considered to be associated with each other if they appear at the same location, during a given night, within some time window of each other. SRI for two mice *i* and *j* is calculated as

$SRI= \frac{X}{X+Y_{i}+Y_{j}+Y_{ij}}$,

where *X* is the number of night-location pairs in which the mice overlap at least once during the night, *Y_ij_* the number of night-location pairs in which both mice appear but not within the overlap window, and *Y_i_* and *Y_j_* the number of night-location pairs in which only mouse *i* or only mouse *j* appears, respectively. SRI is used in a variety of animal behavior studies and allows us to describe spatiotemporal association patterns among the mice (Raulo et al. 2021, Farine and Whitehead 2015). We calculated social networks for the six different groups – three strains, two blocks.

Two major nuances were involved in the calculation of our networks. First, because some mice lost their RFID tags, they did not produce check-in data for periods in which those mice were still active and potentially associating with other mice. To accommodate these missing data, we implemented an adjustment to SRI calculations for these mice, which we referred to as “time control,” again derived from Raulo et al. (2021). For these mice, when determining associations for a pair, we exclude all check-ins taking place after the last night on which the mouse that lost its RFID appeared. This is because the mouse’s absence on those nights could plausibly be attributed to loss of its RFID tag, so we are unable to determine whether its partner in the pair was or was not associating with the mouse with the missing tag. To aid this, we identified associations in two discrete chunks, one prior to the midpoint trapping session, and one after the midpoint trapping session, since the midpoint was the time when missing RFID tags were replaced. With this approach we effectively assume that associations during the phase when the mouse does not have an RFID tag are approximately equal to those occurring at other times of the experiment. Some support for this approach can be found in the fact that the strengths of the pairwise associations between pairs of individuals prior to the midpoint trapping session are well-correlated with the strengths of the association after the midpoint trapping session for that pair (Pearson’s *r* = 0.73; Fig. S7).

The other nuance was in our handling of calculating association strengths for individuals that escaped from their assigned enclosure at some point during the experiment. In these cases, mice would have been physically unable to associate with others in their assigned enclosure. We decided not to excise days of escape from the calculations of association strength for pairs with at least one mouse that escaped from its focal enclosure, since the separation was a genuine factor that altered their frequency of association. Unlike with the lost RFIDs, we do definitively know that the mice of the pair were not associating when one was outside the home enclosure. We did not include escapee mice in the networks of the wedges they entered, however. Thus all of our associations are between mice that shared enclosures.

To explore how location and overlap window duration affected estimates of association strength, we calculated networks in several different scenarios. We used four different overlap window lengths: four hours, one hour, fifteen minutes (our default for presented results), and two minutes. These impose greater or lesser stringency in estimates of association; shorter time frames should be more closely tied to rates of actual contact, simply because associations with these time windows require the mice to be in the same place at a closer time. We also calculated social networks for subsets of readers: just association at the feeding station in each wedge and association at the non-feeding stations in each wedge. These allowed us to assess how association strengths varied at different locations. In general, association strengths at different time intervals were correlated with each other, as were association strengths at different sets of locations, but strengths varied based on the length of the overlap windows and the locations included.

Statistical analyses

Our statistical analyses can be broken down into two major subsets: 1) analyses of the relationships between individual immune traits and individual behavior traits and 2) analyses of the relationship between pairwise immune similarity and social association and other predictors. For all of our statistical analyses, we used a Bayesian framework in R v4.1.2 and the package *brms* (Bürkner 2017, Bürkner et al. 2022). Model selection for each analysis was carried out using widely applicable information criterion (WAIC), a generalized version of Akaike information criterion that can be used with our Bayesian models. The model chosen and presented is that with the lowest WAIC value, except where otherwise specified.

Analyses of individual behavior used mixed effects linear regression models with appropriate distributions for response variables, as follows. Average check-ins per day and average minimum distance traveled were both log-transformed and modeled as normally-distributed response variables. Mean roaming entropy was modeled as a beta-distributed response variable with a logit link function. Immune traits were log-transformed and modeled as normally-distributed response variables. For immune trait values of 0 that could not be log-transformed (e.g. basophil concentrations in the CBC), we assigned values immediately below the detection threshold of the assay to ensure that low values for those traits were not systematically excluded from our analyses. Potential fixed-effect variables included for model selection were genotype and infection status, as well as behavioral traits of interest when investigating the effect of behavior on immune phenotypes. Potential random-effect variables were block of the experiment, enclosure in which the mouse was housed, parentage of the mouse, and the cage in which the mouse was housed prior to release outdoors. For all analyses involving individual behavior either as predictor or response variable, we excluded any mice that either lost an RFID tag or escaped their assigned wedge during the experiment. For analyses of factors predicting pairwise social behavior, we generally used binomial regressions, with the number of trials for each pair of mice being the number of night-location pairs for which at least one of the two mice in the pair appeared and the number of successes being the number of night-location pairs for which the two mice appeared within the designated overlap window.

We took multiple approaches to describing immune similarity as befitting different data. For cell type abundances, we employed methods similar to those used to describe similarity of microbiome composition or species assemblages in community ecology. We assessed the relative abundance of different types of immune cells and calculated the similarity of these abundance distributions as Jaccard index using the R package *vegan* v2.5-7 (Oksanen et al. 2022). Jaccard index is on the interval [0,1], where a value of 0 indicates no overlap at all in the abundance distribution (i.e. entirely dissimilar) and a value of 1 indicates perfect congruence of the abundance distribution (i.e. identical). We calculated similarities in this manner for several different sets of cell types: white blood cell phenotypes from CBC data, combined relative abundances of both CD4 and CD8 T cell memory phenotypes (based on cell type identity determined from CD62L and CD44 expression on T cells as described above) in each of the peripheral blood and mesenteric lymph nodes, as well as just CD4 T cell memory and just CD8 T cell memory in the mesenteric lymph nodes, and relative abundance of different B cell phenotypes on the CD62L/CD44 expression axes in the blood and mesenteric lymph nodes.

For calculating similarity of cytokine phenotypes, we used Manhattan distance, a metric that determines the cumulative difference along each axis between two phenotypes plotted in an *n*-dimensional space, where *n* is the number of traits measured. For plasma cytokines, we first log-transformed the cytokine concentrations so that they were normally-distributed, then scaled the resulting values as Z-scores, such that Manhattan distance would be the sum of the standard deviations of difference between the two phenotypes across all cytokines. For MLN cytokine production, we adjusted production for null controls as stated above, then followed the same procedure.

The analysis of social associations drawn from observation data is considerably more complex and has been the subject of much discussion (Farine 2017, Franks et al. 2021, Puga-Gonzalez et al. 2021, Weiss et al. 2021, Farine and Carter 2022, Hart et al. 2022). We opted to use Bayesian multiple-membership mixed-effects linear regressions, which are recognized as a valid statistical approach when analyzing such relationships (Franks et al. 2021, Weiss et al. 2021) and for which there are statistical packages capable of handling beta regressions (see below). As above, we used model selection to identify the variables for inclusion in our analyses. Because the response variables were describing the status of a pair, all explanatory variables were chosen to match. Potential fixed-effect variables included were the strain identity of the two members of the pair (with the members of the dyad always of the same strain), whether the individuals were full siblings, and whether the individuals were housed in the same cage prior to release in their enclosure; social association was also included as a fixed effect in appropriate cases when analyzing immune similarity. We also included infection status, but we used a different treatment of this variable depending on whether we were analyzing social association or immune similarity. When analyzing social association, we used three categories: both mice infected, one mouse infected, or neither mouse infected. When analyzing immune similarity, we used two categories: shared infection status (meaning either both infected or neither) or different infection status. Potential random-effect variables considered included block of the experiment and the enclosure in which the pair were housed, as well as multiple-membership random effects for each individual in the pair. In general, block, enclosure, and shared parentage were not included in optimal models after model selection, but there were exceptions. Social association levels were modeled as binomially-distributed response variables, where the number of reader-night pairs in which one or both mice appeared (i.e. the denominator of SRI for that pair) was the number of trials and the number of reader-night pairs in which both mice appeared (i.e. the numerator for SRI of that pair) was the number of successes. Jaccard index phenotype similarity variables were modeled as beta-distributed response variables with a logit link function. Beta distributions do not contain 0 or 1, whereas it is possible for our similarity scores to be 0 or 1. However, this is vanishingly unlikely to occur (requiring, respectively, that there be no immune cell types in common or the exact same percentages for each cell type across samples with very large numbers of cells), and we thus considered the use of a beta-distribution as an appropriate choice instead of a zero-and-one-inflated beta regression (Douma and Weedon 2019). Manhattan distance phenotype similarity variables were log-transformed and modeled as normally-distributed response variables. Manhattan distance increases as the similarity of phenotypes decreases; for ease of understanding, we report all coefficient values for regressions predicting Manhattan distance as their negatives, such that a positive coefficient indicates a correlation with increased similarity for both cell distribution similarity and cytokine production similarity.

**Supplemental Text**

Activity levels across the experiment

The number of check-ins in each enclosure varied from day to day (Fig. S8). Because we do not have data from the first few days after release for Block 2, we are limited in our ability to detect trends in changes with activity through time. However, we generally found that the night with peak level of activity for all strains in all enclosures was in the first half of the experiment. Check-in abundance was typically lower in the later part of the experiment, after the mice had been in their enclosures for several weeks. There was one notable exception: in Block 1, the 129S1 mice had few check-ins per night in the first part of the experiment (Fig. S8A). It was only after the midpoint trapping session that check-in numbers per night increased. This disparity is largely in visits to the feeding station in the enclosure (158 pre-midpoint vs. 3,057 post-midpoint), with also a smaller disparity in visits to the reader at the tip of the enclosure. We do not see a similar disparity in the Block 2 129S1 mice or for the Block 2 C57BL/6 mice, which occupied the same enclosure (Enclosure 4) as the Block 1 129S1 mice (Fig. S8B).

Activity levels across the day

The number of check-ins taking place during each hour of the day suggest that the mice were largely nocturnal in their habits. Activity was greatest in the 8 PM–6 AM period (2000–0600) for the 129S1 and C57BL/6 mice (Fig. S9A). The activity distributions by the PWK/PhJ mice were more complex. In Block 1, the PWK/PhJ mice were most active during the night, like the 129S1 and C57BL/6 mice, although activity at RFID readers was more evenly spread compared to the activities of 129S1 and C57BL/6 mice (Fig. S9B). Conversely, in Block 2, the PWK/PhJ mice were active at all times of the day, with a peak in activity during the 6–7 PM hour (1800–1900) (Fig. S9B). However, this peak is driven largely by a single mouse, 307, which had 1,997 check-ins during that hour across the whole experiment. The majority of those 1,997 check-ins came on just two days, 28/7 (776, all in the reader on the left side of the enclosure – 1 check-in every 4.64 seconds) and 8/8 (398, again all in the reader on the left side). Mouse 360 had an unusual peak in activity in the 5–6 PM hour, again with most of the check-ins in that hour coming on just two days, 21/7 (313, all at the tip of the enclosure) and 24/7 (420, all on the left side). These large numbers of check-ins in short time intervals may indicate that the mouse largely stayed inside an RFID reader’s tube during that time, going back and forth and repeatedly passing through the antenna or even sitting within the antenna’s loop. Regardless, even accounting for the mice with unusual behavioral profiles, activity for the Block 2 PWK/PhJ mice was spread fairly evenly throughout the day.

Locations visited

The clear majority of check-ins took place at the feeding stations in each enclosure, and for most individuals the majority of their check-ins took place at the feeding station (Fig. 1C). The PWK/PhJ mice had proportionally more check-ins taking place at the other locations inside their enclosures outside the feeder, followed by C57BL/6 mice, then the 129S1 mice; however, there were not necessarily differences among the strains in an individual’s proportion of check-ins taking place at the feeder (Fig. 1C, Table S1), even if aggregates across all mice differ. There was variation for each mouse strain between blocks in the relative abundances of visits to the different locations – for example, in Block 1 C57BL/6 mice visited the RFID reader at the base of the enclosure only 15 times, while in Block 2 the C57BL/6 mice visited the RFID reader at the base of the enclosure 1,234 times (Fig. S10). And rates of visits to different locations also differed by strain, but the non-feeder locations generally drew visits at similar rates within each strain, with perhaps the reader on the right side of each enclosure having the lowest rate of visits (Table S17).

Roaming entropy

Roaming entropy is a metric that assesses how evenly spread spatially and temporally an individual’s activity is. We calculated roaming entropy for each mouse during each night (8 PM–8 AM), reflecting the hours of greatest activity by the mice. We found that roaming entropy varied significantly within each strain, but 129S1 mice generally had lower roaming entropy than did C57BL/6 or PWK/PhJ mice (Figure S11, Table S18). This would indicate that 129S1 mice were more likely to confine their activity to one RFID reader. Infection with *T. muris* may have also had a weak negative effect on roaming entropy, although further data would be necessary to estimate that relationship with more confidence (Table S18). This similarly accords with the increase in proportion of feeder check-ins associated with infection (Table S1).

Social association under different overlap windows

Levels of social association between mice did vary based on the length of the overlap window used to determine associations. Within each strain, the longer the overlap window, the stronger the average association between pairs of mice (Fig. S1). We further found that the differences between the strains in terms of social association strengths were consistent across the different time windows, and the effects of other variables like infection status and sibling relationships were similarly consistent (Table S2). And for each pair the correlations between social association metrics for different overlap windows were strong, indicating that the relative strengths of pairs’ associations were largely preserved across the different time windows (Table S19). The correlations between association from two-minute windows and association from other windows are the lowest, although still large, perhaps because of the relatively smaller values and smaller windows lead to more random noise in the social association measures.

Individual variation in social association

We assessed the level of association for individuals by determining the average strength of their social associations. This is slightly different from the common metric of degree, the number of individuals that an individual interacts with in a day, because it integrates the number of locations that individuals interact at. In keeping with the pairwise associations, 129S1 mice generally had lower levels of sociality than did C57BL/6 and PWK/PhJ mice, but we found that within a given enclosure the average strength of an individual’s social associations varied relatively little (Fig. S3, Table S20). This means that although individual pairs are variable in their level of social association, individuals cumulatively have similar strengths of social ties. This lack of variation in individual sociality makes it difficult for us to assess any ties between that sociality and immune phenotype and explore the hypotheses in this area.

Just as there were differences between the blocks in levels of individual activity, we can also see differences in individual sociality and pairwise sociality between the blocks, and these differed among the strains. Individual socialities were generally higher for C57BL/6 and PWK/PhJ mice in Block 1 than in Block 2, while the reverse was true for the 129S1 mice. As previously mentioned, the 129S1 mice had relatively few check-ins during the first portion of Block 1, which may partially explain why we see this result. It is possible that these results could be due to the enclosures themselves, with certain enclosures, for whatever reason, more conducive to activity or social association, but we have not identified any consistent activity patterns among the enclosures that might indicate this phenomenon.

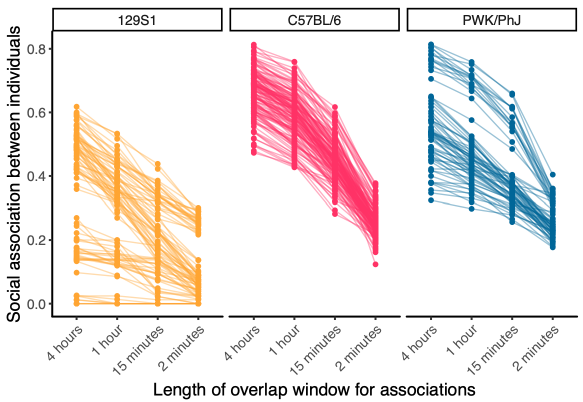

**Figure S1:** Variation in social association for pairs of rewilded mice by length of overlap window for defining association. Pairs including mice that either lost an RFID tag during the experiment or escaped from their home wedge are excluded.

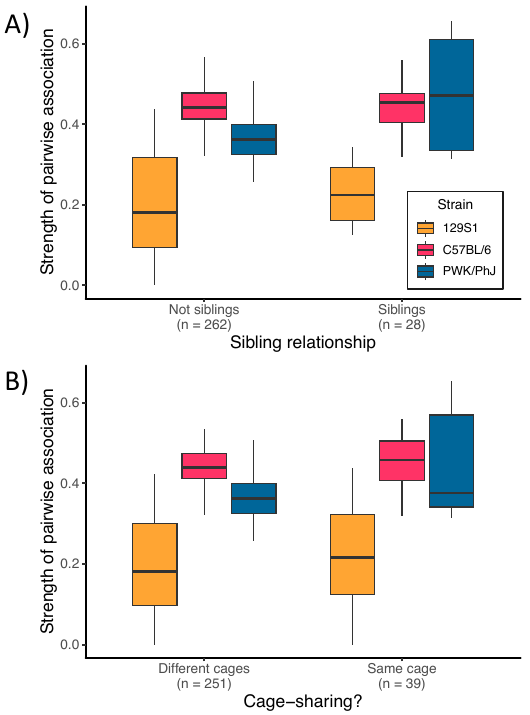

**Figure S2:** Impacts of sibling relationships and cage-sharing on pairwise social association strengths during experiment, broken down by strain. Social associations calculated from 15-minute overlap windows. All pairs containing mice that either escaped during the experiment or lost an RFID tag are excluded. Most pairs that were full siblings also shared cages prior to release. A) Effect of sibling relationships on strength of pairwise associations. B) Effect of cage-sharing prior to release in enclosures on strength of pairwise associations.

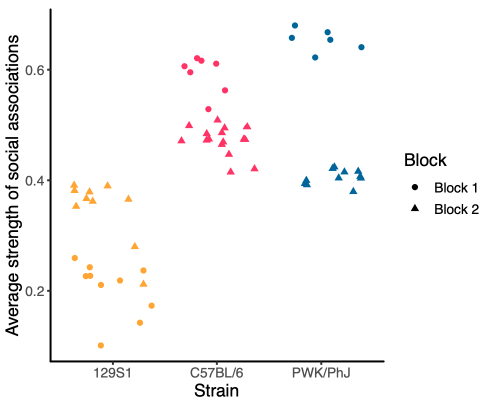

**Figure S3:** Average strength of an individual’s social ties, broken down by strain and block of the experiment. Average strength of social associations is calculated from an average of the pairwise association strengths for all pairs involving that individual. Overlap window length used for defining associations was 15 minutes. Pairs involving individuals with escapes or lost RFID tags were excluded from calculations.

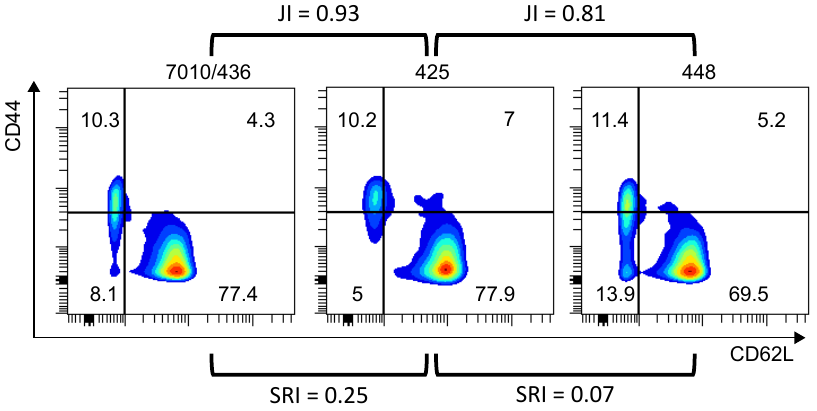

**Figure S4:** Example primary FACS data for two pairs of 129S1 mice from Block 1. Plots show expression of CD62L and CD44 on CD4+ T cells in the mesenteric lymph nodes (MLNs). Horizontal and vertical lines indicate dividing lines for sorting into four memory compartments with unsupervised clustering via Joe’s Flow (see Methods). JI is the Jaccard index for similarity of MLN CD4 T cells; SRI is the social association of the pair (calculated as simple ratio index; see Methods. None of the mice were siblings or shared cages prior to release; 425 and 448 received *T. muris* challenges, while 436 did not.

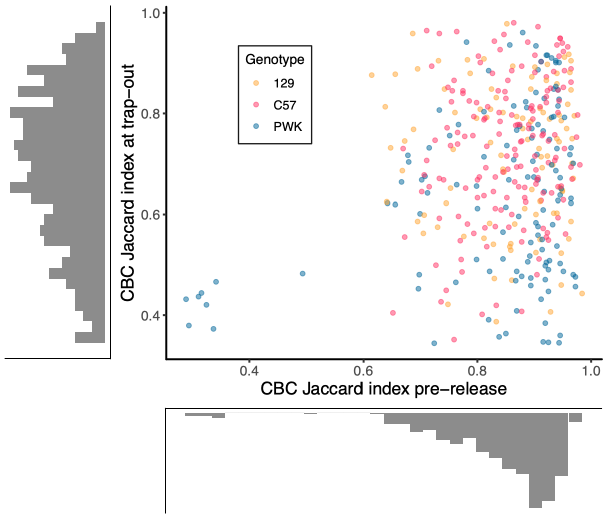

**Figure S5:** Variation among pairs of mice in peripheral blood complete blood count (CBC) immune cell composition prior to release and at the end of the experiment. Pairwise comparisons are only plotted for those mice that were in the same enclosure at the same time (i.e. mice who could have come into contact with each other) and for which CBC data are available both prior to release and at the end of the experiment. Histograms along each axis show distributions of values.

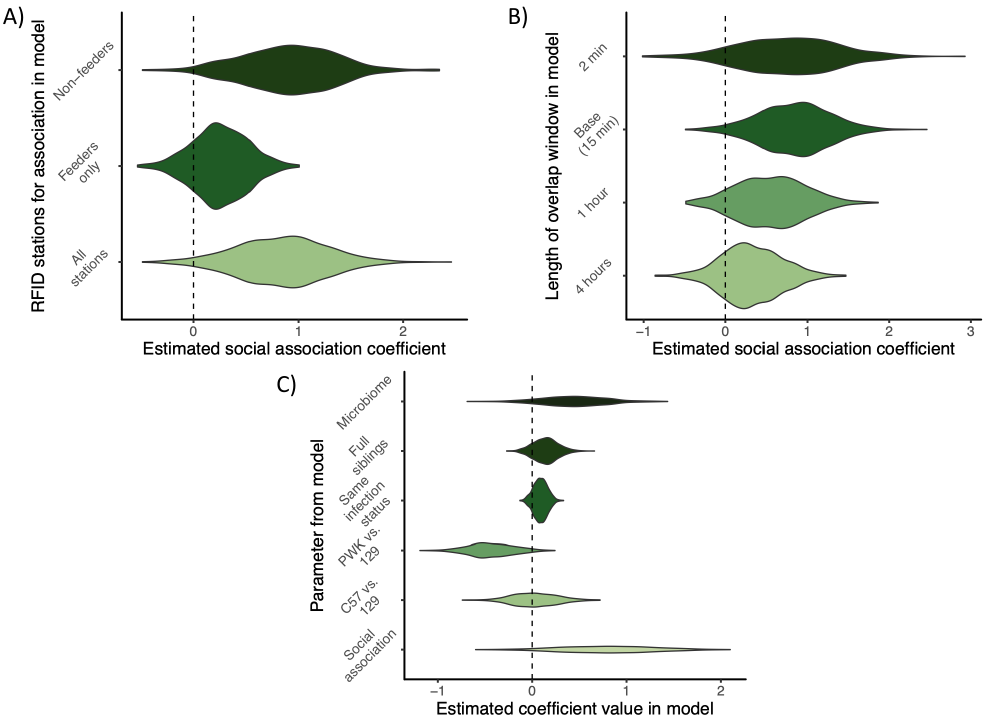

**Figure S6:** Different models of CBC immune similarity and social association to investigate nature of link. Violin plots show regression model coefficient posterior probability distributions plotted via 1000 samples from model-estimated parameter value distribution. A) Estimated relationships between social association calculated from different subsets of RFID readers and similarity of complete blood count (CBC) white blood cell phenotypes. B) Estimated relationships between social association calculated from different overlap windows and CBC phenotype similarity. C) Full results for fixed-effect predictors from model of CBC phenotype similarity in a model that includes gut microbiome similarity. Note that model selection methods do not prefer a model with microbiome similarity.

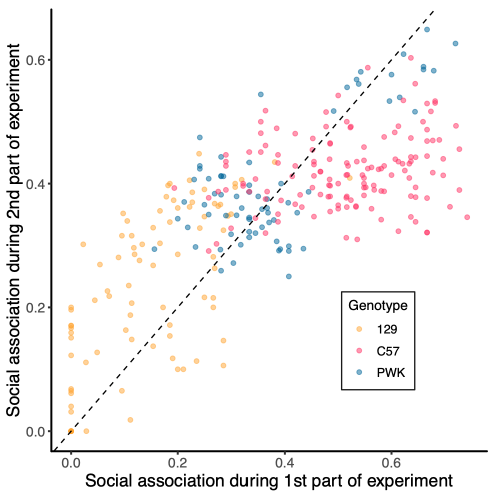

**Figure S7:** Correlation between each mouse pair’s social association prior to midpoint trapping (~2 weeks post-release) and social association after midpoint trapping (from ~2 weeks to trapout ~5 weeks post-release). Dashed line is the line *y* = *x*, where social association between a pair of mice would be identical during the two portions of the experiment. Points above the line are those pairs of mice which associated more after the midpoint trapping; points below the line are those pairs of mice which associated more before the midpoint trapping. Pearson’s *r* = 0.73. All pairs featuring mice that either escaped from their home enclosure or lost and RFID tag during experiment are excluded.

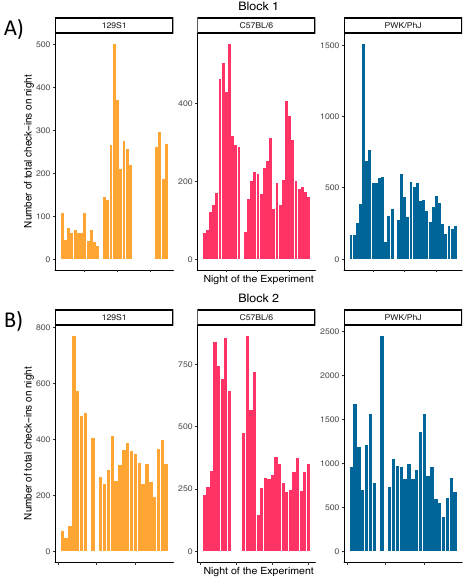

**Figure S8:** Cumulative check-ins across the experiment by day for rewilded mice. Note that since check-in data logging for Block 2 did not start until several days after the mice were released, corresponding columns on the x-axis do not indicate the same time since release in the two Blocks. Nights with no check-ins in an enclosure, due to either a reader malfunction or a trapping session, are nights that were removed from the dataset. In addition, all individuals with lost RFIDs or escapes from enclosures are removed to ensure an unbiased behavioral dataset. Different strains and different blocks contained different numbers of experimental days, which would contribute to differences in cumulative numbers of check-ins. A) Check-ins broken down by strain. B) Check-ins for PWK/PhJ mice broken down by experimental block.

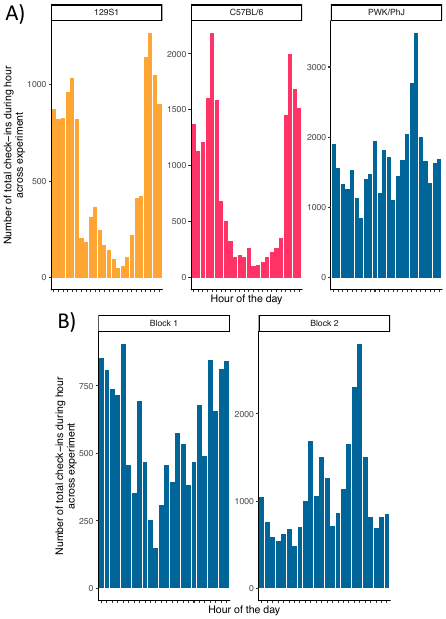

**Figure S9:** Cumulative check-ins across the experiment by hour for rewilded mice, starting from 0000 (12 AM) and running to 2300 (11 PM). All individuals with lost RFIDs or escapes from enclosures are removed to ensure an unbiased behavioral dataset. Different strains and different blocks contained different numbers of experimental days, which would contribute to differences in cumulative numbers of check-ins. A) Check-ins broken down by strain, cumulative across both blocks. B) Check-ins for PWK/PhJ mice broken down by experimental block.

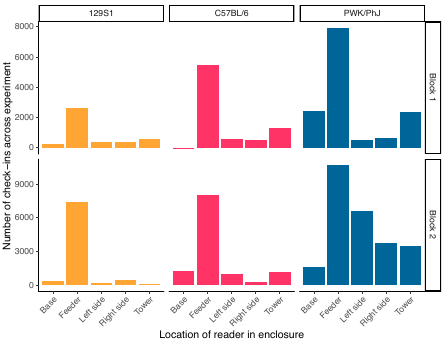

**Figure S10:** Cumulative check-ins across the experiment by location for the rewilded mice, broken down by strain and experimental block. All individuals with lost RFIDs or escapes from enclosures are removed to ensure an unbiased behavioral dataset. Different strains and different blocks had different numbers of individuals and experimental days, which would contribute to differences in cumulative numbers of check-ins.

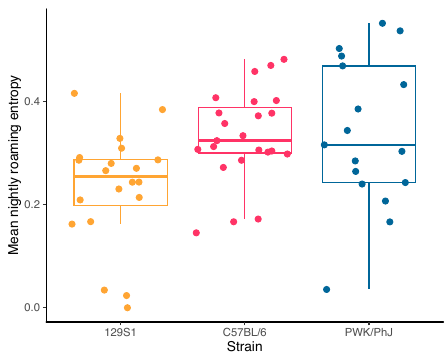

**Figure S11:** Mean nightly roaming entropy for rewilded mice, broken down by strain. Roaming entropy is calculated only for the 8 PM–8 AM hours, as detailed in the methods. All individuals with lost RFIDs or escapes from enclosures are removed to ensure an unbiased behavioral dataset.

**Table S1:** Model results from an analysis of abundance of RFID check-ins per night taking place across the whole experiment for each mouse. Model is a Bayesian linear regression with log-transformed number of check-ins per night as the normally-distributed response variable. Predictors determined after model selection; fixed effects were mouse strain and mouse infection status, while no random effects were included.

| Parameter | Mean model estimate | 95% highest posterior density interval |
| --- | --- | --- |
| C57BL/6 vs. 129S1 | 0.24 | [0.02, 0.45] |
| PWK/PhJ vs. 129S1 | 0.66 | [0.43, 0.89] |
| Infected vs. uninfected | -0.03 | [-0.22, 0.17] |

**Table S2:** Model results from an analysis of abundance of RFID check-ins per night taking place across the whole experiment for each mouse. Model is a Bayesian linear regression with log-transformed number of check-ins per night as the normally-distributed response variable. Predictors determined after model selection; the fixed effects were mouse strain and block, while random effects were enclosure and parentage.

| Parameter | Mean model estimate | 95% highest posterior density interval |
| --- | --- | --- |
| C57BL/6 vs. 129S1 | 0.49 | [0.16, 0.79] |
| PWK/PhJ vs. 129S1 | 0.61 | [0.30, 0.89] |
| Block 2 vs. Block 1 | -0.12 | [-0.34, 0.10] |

**Table S3:** Model results from an analysis of proportion of RFID check-ins taking place at feeder stations for each mouse. Model is a Bayesian linear regression with binomial response variable, with overall number of check-ins being the trials and number of feeder check-ins being the successes. Predictors determined after model selection; random effects were mouse parentage and mouse cage prior to release. Models without mouse strain were strongly preferred by WAIC (∆WAIC = -15.2) and produced similar estimates for the effect of infection status.

| Parameter | Mean model estimate | 95% highest posterior density interval |
| --- | --- | --- |
| **C57BL/6 vs. 129S1** | 0.09 | [-1.09, 1.18] |
| **PWK/PhJ vs. 129S1** | -0.69 | [-1.92, 0.46] |
| **Infected vs. uninfected** | 0.52 | [0.47, 0.57] |

**Table S4:** Model results from an analysis of social association between pairs of mice. Model is a Bayesian binomial regression. The first model (“Whole experiment”) looks at predictors for social association across the whole of the experiment; the second (“Post-infection”) looks at predictors for social association during only the portion of the experiment after challenges with *Trichuris muris* eggs were conducted. For both of these models the overlap window used is 15 minutes; the other three use different overlap windows. Values in table are mean parameter estimates and, in brackets, boundaries of 95% highest posterior density intervals. Predictors determined after model selection; random effects included individual ID and, only in the post-infection model, both block of experiment and enclosure (“wedge”).

| **Parameter** | **Whole experiment** | **Post-infection** | **Whole w/4-hour overlap** | **Whole w/1-hour overlap** | **Whole w/2-minute overlap** |
| --- | --- | --- | --- | --- | --- |
| **C57BL/6 vs. 129S1** | 1.52  [0.87, 2.10] | 1.48  [0.85, 2.12] | 1.49  [0.65, 2.22] | 1.58  [0.87, 2.30] | 1.36  [0.80, 1.93] |
| **PWK/PhJ vs. 129S1** | 1.40  [0.80, 1.98] | 1.16  [0.53, 1.80] | 1.20  [0.38, 2.00] | 1.33  [0.56, 2.13] | 1.41  [0.86, 1.93] |
| **Neither infected vs. both infected in dyad** | 0.02  [-0.48, 0.48] | 0.12  [-0.36, 0.58] | 0.01  [-0.67, 0.72] | 0.03  [-0.57, 0.69] | 0.13  [-0.34, 0.60] |
| **One infected vs. both infected in dyad** | -0.02  [-0.27, 0.22] | 0.03  [-0.21, 0.27] | -0.03  [-0.39, 0.32] | -0.03  [-0.33, 0.30] | 0.04  [-0.20, 0.28] |
| **Cage-sharing pre-release** | 0.06  [-0.05, 0.17] | 0.05  [-0.08, 0.19] | 0.03  [-0.08, 0.15] | 0.02  [-0.09, 0.14] | -0.01  [-014, 0.12] |
| **Full siblings** | -0.02  [-0.16, 0.12] | -0.01  [-0.18, 0.16] | 0.01  [-0.12, 0.14] | 0.01  [-0.12, 0.14] | -0.02  [-0.19, 0.15] |

**Table S5:** Model results from an analysis of number of check-ins per night taking place after *T. muris* challenge for each mouse. Model is a Bayesian linear regression with a normally-distributed response variable – log_10_(check-ins per night). Predictors determined after model selection; no random effects were included.

| **Parameter** | **Mean model estimate** | **95% highest posterior density interval** |
| --- | --- | --- |
| **C57BL/6 vs. 129S1** | 0.00 | [-0.16, 0.15] |
| **PWK/PhJ vs. 129S1** | 0.46 | [0.29, 0.63] |
| **Infected vs. uninfected** | -0.16 | [-0.31, -0.02] |

**Table S6:** Model results from analyses of relationships between metrics of individual behavior and abundances of white blood cell types. All models are Bayesian linear regressions with a normally-distributed response variable – log_10_(cell type abundance). Predictors are chosen via model selection methods; other fixed-effect predictors with results not shown here include strain and block of the experiment. No random effects were included in any of the models. Before use as predictors, check-ins per night and minimum distance traveled per night were both log_10_-transformed. All mice that either lost an RFID tag or escaped from their assigned enclosure during the experiment were excluded (*n* = 60)*.* Numbers presented in table are means and, in brackets, boundaries of 95% highest posterior density intervals for parameter estimates.

| **Cell Type** | **Mean and 95% HPDI range for association with check-ins per night** | **Mean and 95% HPDI range for minimum distance per night** |
| --- | --- | --- |
| **Neutrophils** | -0.02  [-0.62, 0.56] | -0.03  [-0.58, 0.54] |
| **Lymphocytes** | 0.35  [-0.21, 0.90] | 0.36  [-0.13, 0.87] |
| **Eosinophils** | 0.41  [-0.40, 1.22] | -0.28  [-0.92, 0.42] |
| **Basophils** | 0.28  [-0.24, 1.21] | 0.36  [-0.24, 0.93] |
| **Monocytes** | -0.05  [-0.67, 0.60] | -0.56  [-1.07, -0.03] |

**Table S7:** Model results for analyses of pairwise immune similarity, for several aspects of cellular immune phenotype. All models are Bayesian linear regressions with immune similarity as a beta-distributed response variable. Pairwise immune similarities calculated with Jaccard index using relative abundances of different immune cell types (see Materials and Methods). Predictors chosen via model selection methods; ∆WAIC shows difference between WAIC for models of immune similarity with and without social association (positive values indicate a preference for models with social association as a predictor). Individual ID is included as a random effect in all models. Values in table are means and, in brackets, bounds of 95% highest posterior density intervals for parameter estimates.

| **Parameter** | **CBC** | **CD4/CD8 T cells in MLNs** | **CD4 T cells*** | **CD8 T cells*** | **CD4/CD8 T cells in blood** | **B cells in MLNs** |
| --- | --- | --- | --- | --- | --- | --- |
| **Social association** | 0.87  [-0.06, 1.80] | 1.15  [0.37, 1.93] | 1.58  [0.67, 2.52] | 0.74  [-0.40, 1.80] | -0.17  [-0.96, 0.63] | 1.00  [-0.16, 2.18] |
| **C57BL/6 vs. 129S1** | -0.08  [-0.52, 0.41] | -0.58  [-1.46, -0.38] | -0.66  [-1.42, 0.13] | -0.59  [-1.86, 0.55] | -0.24  [-0.62, 0.15] | -0.31  [-0.99, 0.36] |
| **PWK/PhJ vs. 129S1** | -0.47  [-0.92, 0.00] | -1.03  [-1.87, -0.06] | -1.22  [-1.90, -0.49] | -0.86  [-2.05, 0.36] | -0.58  [-0.96, -0.19] | -0.62  [-1.33, 0.08] |
| **Shared infection status** | 0.08  [-0.05, 0.20] | 0.14  [0.06, 0.22] | 0.19  [0.08, 0.31] | 0.05  [-0.06, 0.15] | 0.02  [-0.10, 0.15] | 0.30  [0.15, 0.44] |
| **Full siblings** | 0.14  [-0.07, 0.36] | -0.08  [-0.21, 0.05] | -0.08  [-0.25, 0.10] | -0.05  [-0.23, 0.13] | -0.09  [-0.30, 0.12] | -0.14  [-0.37, 0.09] |
| **∆WAIC** | 1.1 | 1.1 | 2.4 | -2.7 | -0.2 | 0.0 |

**Table S8:** Model results for analyses of pairwise immune similarity, for two aspects of cytokine immune phenotype. All models are Bayesian linear regressions with log-transformed cytokine abundance or production similarity as a normally-distributed response variable. Pairwise plasma cytokine similarity calculated with Manhattan distance from absolute cytokine abundances in blood. MLN cytokine production similarity calculated from log_2_-fold increase in cytokine production after stimulation with antigens with Manhattan distance (see Materials and Methods). Predictors chosen via model selection methods; ∆WAIC shows difference between WAIC for models of immune similarity with and without social association (positive values indicate a preference for models with social association). Individual ID is included as a random effect in all models. Values in table are means and, in brackets, bounds of 95% highest posterior density intervals for parameter estimates.

| **Parameter** | **Plasma cytokines** | **MLN cytokine production** |
| --- | --- | --- |
| **Social association** | -0.25  [-0.70, 0.20] | -1.09  [-1.72, -0.45] |
| **C57BL/6 vs. 129S1** | 0.08  [-0.20, 0.34] | 0.44  [0.06, 0.87] |
| **PWK/PhJ vs. 129S1** | 0.31  [-0.06, 0.53] | 0.21  [-0.19, 0.63] |
| **Shared infection status** | 0.15  [-0.08, 0.23] | 0.16  [0.08, 0.24] |
| **Full siblings** | 0.08  [-0.03, 0.20] | 0.06  [-0.05, 0.16] |
| **∆WAIC** | 0.3 | 7.0 |

**Table S9:** Model results for analyses of pairwise CBC immune similarity at several different times during the experiment. All models are Bayesian linear regressions with immune similarity as a beta-distributed response variable. Pairwise immune similarities calculated with Jaccard index using relative abundances of different immune cell types (see Materials and Methods). All blood samples were acquired via cheek bleeds. Starting blood samples were taken prior to release, midpoint blood samples were taken two weeks after release, prior to any challenge with *T. muris*, and trapout blood samples were taken prior to sacrifice. Predictors chosen via model selection methods; individual ID is included as a random effect in all models. Values in table are means and, in brackets, bounds of 95% highest posterior density intervals for parameter estimates.

| **Parameter** | **Start** | **Midpoint** | **Trapout** |
| --- | --- | --- | --- |
| **Social association** | 0.12  [-0.45, 0.73] | 0.30  [-0.33, 0.98] | 0.87  [-0.06, 1.80] |
| **C57BL/6 vs. 129S1** | 0.02  [-0.46, 0.48] | -0.09  [-0.52, 0.34] | -0.08  [-0.52, 0.41] |
| **PWK/PhJ vs. 129S1** | -0.01  [-0.45, 0.43] | 0.12  [-0.31, 0.51] | -0.47  [-0.92, 0.00] |
| **Shared infection status** | -0.08  [-0.17, 0.02] | 0.05  [-0.08, 0.18] | 0.08  [-0.05, 0.20] |
| **Full siblings** | 0.15  [-0.01, 0.33] | 0.07  [-0.13, 0.26] | 0.14  [-0.07, 0.36] |

**Table S10:** Model results for analyses of pairwise MLN CD4 T cell memory similarity using pairwise social association calculated for different subsets of RFID reader location. All models are Bayesian linear regressions with immune similarity as a beta-distributed response variable. Pairwise immune similarities calculated with Jaccard index using relative abundances of different immune cell types (see Materials and Methods). Predictors chosen via model selection methods; individual ID is included as a random effect in all models. Values in table are means and, in brackets, bounds of 95% highest posterior density intervals for parameter estimates.

| **Parameter** | **Base model** | **Feeders only** | **Non-feeders only** |
| --- | --- | --- | --- |
| **Social association** | 1.58  [0.67, 2.52] | 1.25  [0.58, 1.91] | 1.20  [0.32, 2.12] |
| **C57BL/6 vs. 129S1** | -0.66  [-1.42, 0.13] | -0.63  [-1.35, 0.10] | -0.52  [-1.20, 0.15] |
| **PWK/PhJ vs. 129S1** | -1.22  [-1.90, -0.49] | -1.31  [-2.02, -0.57] | -1.12  [-1.91, -0.32] |
| **Shared infection status** | 0.19  [0.08, 0.31] | 0.21  [0.10, 0.33] | 0.07  [0.19, 0.31] |
| **Full siblings** | -0.08  [-0.25, 0.10] | -0.06  [-0.24, 0.13] | -0.08  [-0.25, 0.10] |

**Table S11:** Model results for analyses of pairwise CBC immune similarity using pairwise social association calculated for different subsets of RFID reader location. All models are Bayesian linear regressions with immune similarity as a beta-distributed response variable. Pairwise immune similarities calculated with Jaccard index using relative abundances of different immune cell types (see Materials and Methods). Predictors chosen via model selection methods; individual ID is included as a random effect in all models. Values in table are means and, in brackets, bounds of 95% highest posterior density intervals for parameter estimates.

| **Parameter** | **Base model** | **Feeders only** | **Non-feeders only** |
| --- | --- | --- | --- |
| **Social association** | 0.87  [-0.06, 1.80] | 0.27  [-0.22, 0.77] | 0.91  [0.02, 1.80] |
| **C57BL/6 vs. 129S1** | -0.08  [-0.52, 0.41] | 0.04  [-0.45, 0.52] | -0.06  [-0.54, 0.42] |
| **PWK/PhJ vs. 129S1** | -0.47  [-0.92, 0.00] | -0.42  [-0.92, 0.07] | -0.44  [-0.94, -0.04] |
| **Shared infection status** | 0.08  [-0.05, 0.20] | 0.08  [-0.04, 0.21] | 0.08  [-0.04, 0.20] |
| **Full siblings** | 0.14  [-0.07, 0.36] | 0.14  [-0.07, 0.37] | 0.13  [-0.08, 0.35] |

**Table S12:** A table showing negative serology and PCR results for common mouse pathogens (see attached spreadsheet – formatting constraints make it difficult to fit in a standard document).

**Table S13:** Model results for analyses of pairwise immune similarity, for several aspects of cellular immune phenotype, incorporating pairwise bacterial gut microbiome similarity. All models are Bayesian linear regressions with immune similarity as a beta-distributed response variable. Pairwise immune similarities calculated with Jaccard index using relative abundances of different immune cell types (see Materials and Methods). Pairwise microbiome similarity calculated from microbiome composition as assessed with 16S sequencing of DNA extracted from fecal pellets. Predictors chosen via model selection methods; ∆WAIC shows difference between WAIC for models of immune similarity with and without social association (positive values indicate a preference for models with microbiome). Individual ID is included as a random effect in all models. Values in table are means and, in brackets, bounds of 95% highest posterior density intervals for parameter estimates.

| **Parameter** | **CD4/CD8 T cells in MLNs** | **CD4 T cells in MLNs** | **CBC** | **B cells** | **Microbiome** |
| --- | --- | --- | --- | --- | --- |
| **Social association** | 0.77  [-0.04, 1.66] | 1.18  [0.14, 2.25] | 0.78  [-0.14, 1.68] | 0.52  [-0.79, 1.90] | -0.56  [-1.33, 0.19] |
| **C57BL/6 vs. 129S1** | -0.57  [-1.49, 0.31] | -0.54  [-1.34, 0.28] | 0.03  [-0.40, 0.47] | -0.21  [-0.97, 0.52] | 0.35  [-0.38, 1.09] |
| **PWK/PhJ vs. 129S1** | -0.57  [-1.43, 0.26] | -0.81  [-1.56, -0.01] | -0.45  [-0.87, -0.01] | -0.28  [-0.98, 0.41] | 0.67  [-0.04, 1.38] |
| **Shared infection status** | 0.17  [0.07, 0.26] | 0.26  [0.12, 0.38] | 0.09  [-0.05, 0.23] | 0.26  [0.09, 0.44] | -0.00  [-0.09, 0.09] |
| **Full siblings** | -0.04  [-0.20, 0.12] | 0.00  [-0.21, 0.23] | 0.14  [-0.11,0.40] | -0.14  [-0.41, 0.15] | 0.07  [-0.08, 0.23] |
| **Microbiome similarity** | -0.15  [-0.70, 0.43] | -0.40  [-1.10, 0.33] | 0.47  [-0.18, 1.11] | -0.27  [-1.12, 0.58] | N/A |
| **∆WAIC** | -1.5 | -1.1 | -2.8 | -0.6 | N/A |

**Table S14:** Model results for analyses of pairwise MLN CD4 T cell memory similarity using pairwise social association calculated with different length overlap windows for determining association. All models are Bayesian linear regressions with immune similarity as a beta-distributed response variable. Pairwise immune similarities calculated with Jaccard index using relative abundances of different immune cell types (see Materials and Methods). Predictors chosen via model selection methods; individual ID is included as a random effect in all models. Values in table are means and, in brackets, bounds of 95% highest posterior density intervals for parameter estimates.

| **Parameter** | **4 hours** | **1 hour** | **15 minutes** | **2 minutes** |
| --- | --- | --- | --- | --- |
| **Social association** | 1.23  [0.36, 2.09] | 1.50  [0.61, 2.34] | 1.58  [0.67, 2.52] | 0.95  [-0.19, 2.07] |
| **C57** | -0.63  [-1.40, 0.10] | -0.69  [-1.43, 0.05] | -0.66  [-1.42, 0.13] | -0.36  [-1.04, 0.35] |
| **PWK** | -1.09  [-1.79, -0.36] | -1.22  [-1.96, -0.50] | -1.22  [-1.90, -0.49] | -1.08  [-1.84, -0.34] |
| **Same infection status** | 0.19  [0.07, 0.31] | 0.19  [0.08, 0.31] | 0.19  [0.08, 0.31] | 0.19  [0.07, 0.31] |
| **Full siblings** | -0.08  [-0.26, 0.11] | -0.09  [-0.27, -0.10] | -0.08  [-0.25, 0.10] | -0.07  [-0.25, 0.11] |

**Table S15:** Model results for analyses of pairwise CBC immune similarity using pairwise social association calculated with different length overlap windows for determining association. All models are Bayesian linear regressions with immune similarity as a beta-distributed response variable. Pairwise immune similarities calculated with Jaccard index using relative abundances of different immune cell types (see Materials and Methods). Predictors chosen via model selection methods; individual ID is included as a random effect in all models. Values in table are means and, in brackets, bounds of 95% highest posterior density intervals for parameter estimates.

| **Parameter** | **4 hours** | **1 hour** | **15 minutes** | **2 minutes** |
| --- | --- | --- | --- | --- |
| **Social association** | 0.34  [-0.38, 1.05] | 0.57  [-0.22, 1.33] | 0.87  [-0.06, 1.80] | 0.73  [-0.41, 1.91] |
| **C57BL/6 vs. 129S1** | 0.03  [-0.45, 0.51] | -0.03  [-0.51, 0.49] | -0.08  [-0.52, 0.41] | 0.04  [-0.43, 0.46] |
| **PWK/PhJ vs. 129S1** | -0.40  [-0.87, 0.07] | -0.43  [-0.81, 0.06] | -0.47  [-0.92, 0.00] | -0.41  [-0.89, 0.07] |
| **Shared infection status** | 0.08  [-0.04, 0.20] | 0.08  [-0.04, 0.19] | 0.08  [-0.05, 0.20] | 0.08  [-0.04, 0.20] |
| **Full siblings** | 0.14  [-0.07, 0.36] | 0.14  [-0.09, 0.36] | 0.14  [-0.07, 0.36] | 0.14  [-0.06, 0.36] |

**Table S16:** Fluorescent antibodies used for labeling lymphoid cells via flow cytometry.

| **REAGENT or RESOURCE** | **SOURCE** | **IDENTIFIER** |
| --- | --- | --- |
| Antibodies | | |
| Anti-mouse CD45 (30-F11) BUV395 | BD Biosciences | Cat#: 564279;  RRID: AB_2651134 |
| Rat Anti-mouse CD103 (M290) BUV496 | BD Biosciences | Cat#: 741083;  RRID:AB_2870687 |
| Rat Anti-mouse CD62L (MEL-14) BUV563 | BD Biosciences | Cat#: 741230;  RRID: AB_2870784 |
| Hamster Anti-mouse TCR-B chain (H57-597) BUV737 | BD Biosciences | Cat#: 612821 |
| Rat Anti-Mouse CD44 (IM7) BUV805 | BD Biosciences | Cat#: 741921  RRID: AB_2871234 |
| Rat Anti-Mouse NKp46 CD335 (29A1.4), BV480 | BD Biosciences | Cat#: 746264  RRID: AB_2743596 |
| Anti-Mouse NK-1.1 (PK136), BUV661 | BD Biosciences | Cat#:741477  RRID: AB_2870942 |
| Hamster Anti-Mouse TCR β Chain, (H57-597), BUV737 | BD Biosciences | Cat#: 612821 |
| Anti-mouse CD8β (H35-17.2) BV510 | BioLegend | Cat#:103278 RRID: AB_2860603 |
| Anti-mouse CD4 (RM4-5) BV570 | BioLegend | Cat#: 100542 RRID: AB_2563051 |
| Hamster Anti-Mouse KLRG1 (2F1), BV750 | BD Biosciences | Cat#: 746972;  AB_2871754 |
| Anti-mouse TCRγδ (eBio-GL3) PECy5 | Thermofisher | Cat#: **15-5711-82** RRID:  AB_468804 |
| Anti-mouse CD45R/B220 (RA3-6B2) APC/Fire810 | BioLegend | Cat#: 103278 RRID: AB_2860603 |
| Ant-mouse Tbet (4B10) BV421 | Biolegend | Cat#”: 644816  RRID: AB_10959653 |
| Ki-67 Monoclonal Antibody (SolA15) AF532 | Thermofisher | Cat#: 58-5698-82  RRID: AB_2802365 |
| Anti-mouse FoxP3 (FJK-16) AF700 | BioLegend | Cat#:320014 RRID: AB_439750 |
| Anti-mouse RorgT (Q31-378) PE-CF594 | BD Biosciences | Cat# 56284 RRID: AB_2651150 |
| Gata-3 Monoclonal Antibody (TWAJ), Alexa Fluor™ 488 | Thermofisher | Cat #53-9966-42; RRID :AB_2574493 |
| Hamster Anti-Mouse CD69 (H1.2F3) BV650 | BD Biosciences | Cat # 740460; RRID: AB_2740186 |
| Anti-Mouse CD279 (PD-1) (29F.1A12) BV421 | BioLegend | Cat: #135218; RRID: AB_2561447 |
| Anti-mouse CD16/32 Fc Block™ (KT1632) | BD Biosciences | Cat#: **MA5-18012** RRID:  AB_2539396 |

**Table S16:** Results for regression analysis of rates of check-ins per night at each location in the enclosure for mice. We excluded all mice with lost RFIDs or escapes for an unbiased assessment of behavior. Check-ins per night at each location was log_10_-transformed and modeled with a Gaussian distribution. Fixed effects included location, mouse strain, mouse infection status, and the block of the experiment; mouse ID was included as a random effect.

| Parameter | Mean coefficient estimate | 95% highest posterior density interval |
| --- | --- | --- |
| Intercept | 0.09 | [-0.15, 0.30] |
| Feeding station vs. base | 0.98 | [0.84, 1.14] |
| Left side vs. base | -0.05 | [-0.21, 0.11] |
| Right side vs. base | -0.16 | [-0.32, 0.00] |
| Tower reader vs. base | 0.06 | [-0.10, 0.22] |
| C57BL/6 vs. 129S1 | 0.27 | [0.09, 0.46] |
| PWK/PhJ vs. 129S1 | 0.67 | [0.46, 0.88] |
| Infected vs. uninfected | -0.08 | [-0.23, 0.09] |
| Block 1 vs. Block 2 | -0.05 | [-0.121, 0.12] |

**Table S17:** Results for regression analysis of mean roaming entropy per night for rewilded mice. Mean roaming entropy is modeled as a beta-distributed response variable, with strain and infection status as fixed effects and block of the experiment and enclosure as random effects. Cage of origin prior to release and parentage were excluded as random effects after model selection. All individuals with lost RFIDs or escapes from enclosures are removed to ensure an unbiased behavioral dataset.

| Parameter | Mean model estimate | 95% highest posterior density interval |
| --- | --- | --- |
| C57BL/6 vs. 129S1 | 0.70 | [0.28, 1.11] |
| PWK/PhJ vs. 129S1 | 0.74 | [0.27, 1.19] |
| Infected vs. uninfected | -0.22 | [-0.53, 0.08] |

**Table S19:** Pearson’s *r* for correlations between the social association strength for a pair calculated from various overlap windows. All individuals with lost RFIDs or escapes from enclosures are removed to ensure an unbiased behavioral dataset.

|  | 4 hours | 1 hour | 15 minutes | 2 minutes |
| --- | --- | --- | --- | --- |
| **4 hours** | X | 0.98 | 0.92 | 0.78 |
| **1 hour** | 0.98 | X | 0.96 | 0.82 |
| **15 minutes** | 0.92 | 0.96 | X | 0.89 |
| **2 minutes** | 0.78 | 0.82 | 0.89 | X |

**Table S20:** Model results for analyses of average strength of an individual’s pairwise social associations. Pairwise social association is modeled as a beta-distributed response variable and is calculated from an average of the pairwise association strengths for all pairs involving that individual. Individuals with escapes or lost RFID tags were excluded from the analysis (*n* = 60). Predictors were chosen via model selection methods; enclosure was included as a random effect.

| **Parameter** | **Mean model estimate** | **95% highest posterior density interval** |
| --- | --- | --- |
| **C57BL/6 vs. 129S1** | 1.52 | [1.38, 1.65] |
| **PWK/PhJ vs. 129S1** | 1.02 | [0.87, 1.16] |
| **Block 2 vs. Block 1** | -0.25 | [-0.35, -0.15] |
